# Enhanced guide-RNA Design and Targeting Analysis for Precise CRISPR Genome Editing of Single and Consortia of Industrially Relevant and Non-Model Organisms

**DOI:** 10.1101/139626

**Authors:** Brian J. Mendoza, Cong T. Trinh

## Abstract

**Motivation:** Genetic diversity of non-model organisms offers a repertoire of unique phenotypic features for exploration and cultivation for synthetic biology and metabolic engineering applications. To realize this enormous potential, it is critical to have an efficient genome editing tool for rapid strain engineering of these organisms to perform novel programmed functions.

**Results:** To accommodate the use of CRISPR/Cas systems for genome editing across organisms, we have developed a novel method, named CASPER (CRISPR Associated Software for Pathway Engineering and Research), for identifying on- and off-targets with enhanced predictability coupled with an analysis of non-unique (repeated) targets to assist in editing any organism with various endonucleases. Utilizing CASPER, we demonstrated a modest 2.4% and significant 30.2% improvement (F-test, p<0.05) over the conventional methods for predicting on- and off-target activities, respectively. Further we used CASPER to develop novel applications in genome editing: multitargeting analysis (i.e. simultaneous multiple-site modification on a target genome with a sole guide-RNA (gRNA) requirement) and multispecies population analysis (i.e. gRNA design for genome editing across a consortium of organisms). Our analysis on a selection of industrially relevant organisms revealed a number of non-unique target sites associated with genes and transposable elements that can be used as potential sites for multitargeting. The analysis also identified shared and unshared targets that enable genome editing of single or multiple genomes in a consortium of interest. We envision CASPER as a useful platform to enhance the precise CRISPR genome editing for metabolic engineering and synthetic biology applications.

## 1. Introduction

Transforming biology into engineering practice has shaped the frontiers of synthetic biology and metabolic engineering (Connelly, et al., 2015; Trinh and Mendoza, 2016). This helps drive industrialization of biology with broad applications related to food, health, energy, and the environment from production of drugs to fight diseases to synthesis of renewable fuels to replace fossil fuels (Nielsen and Keasling, 2016). Genetic diversity of non-model organisms offers a repertoire of unique phenotypic features for exploration and cultivation for these applications (Jullesson, et al., 2015; Lee, et al., 2012). To realize this enormous potential, it is critical to have an efficient genome editing tool for rapid strain engineering of these organisms to perform novel programmed functions. In the last several years, the CRISPR technology has emerged as a powerful genome editing tool for metabolic engineering and synthetic biology applications (Cong, et al., 2013; Farasat and Salis, 2016; Garst, et al., 2016; Gomaa, et al., 2014; Hsu, et al., 2014; Jakočiūnas, et al., 2015; Li, et al., 2013; Platt, et al., 2014; Qi, et al., 2012; Qi, et al., 2013; Sander and Joung, 2014; Zeitoun, et al., 2015).

Using a CRISPR/Cas system for genome editing requires it to be both highly active and accurate. CRISPR tools that are used to design guide RNAs (gRNAs) to form ribonucleoprotein (RNP) complexes for effective genome targeting focus on two types of evaluation. The first type is the prediction of “on-target” activity of a selected gRNA on a target DNA (or RNA) sequence on the genome. This on-target activity reflects the ability of such a gRNA to successfully find and bind the Cas endonuclease to the target sequence. The second type is the prediction of “off-target” activity of a selected gRNA, which pertains to the propensity of the RNP complex to interact with sequences on the genome similar to the target sequence.

Choosing gRNA sequences with high activity is paramount to all genome editing endeavors for metabolic engineering and synthetic biology applications (Feng, et al., 2015; Ryan, et al., 2014; Stovicek, et al., 2015). To predict which sequences lend themselves to efficient targeting, various algorithms have been developed based on high-throughput experimentation and validation (Doench, et al., 2016; Doench, et al., 2014; Moreno-Mateos, et al., 2015). By identifying prominent nucleotide features in highly active gRNAs and their target sequences, scoring tables were generated and a gRNA sequence may be cross-referenced to predict its on-target activity. Past experimental evidence has revealed that the CRISPR RNP complex can reach cleavage efficiencies of at least 80% for single site modification (Jakočiūnas, et al., 2015). However, this efficiency might become problematic when applying a CRISPR/Cas system to modify multiple sites across the genome simultaneously using multiple gRNAs, known as multiplexing, for rapid strain engineering. For instance, the probability of achieving activity at 5 loci simultaneously drops to 32% (0.85). Therefore, developing new algorithms capable of accurately designing gRNA sequences with high activity is important for multiplex CRISPR implementation.

In addition to the on-target analysis, it is very critical to make sure that a gRNA design has limited activity at non-targeted sites. The foundation for such off-target analysis for the spCas9 (Cas9 derived from *Streptococcus pyogenes*) system in eukaryotes was established by Hsu *et al*. (Hsu, et al., 2013) and later by Lin *et al*. (Lin, et al., 2014). Studies building on this foundation generally take two directions: one being the engineering or discovery of alternative Cas enzymes that exhibit lower off-target activity (Kleinstiver, et al., 2015), and the other being algorithms that enhance prediction capabilities (Lin, et al., 2014). Recently developed software platforms have implemented various algorithms to determine off-target sites for a gRNA design, based on either biological principles or training a model to fit experimental data (Doench, et al., 2016; Hendel, et al., 2015; Kleinstiver, et al., 2016; Labun, et al., 2016). A collection of these programs is detailed at https://omictools.com/crispr-cas9-category. Unfortunately, some potential off-target sequences are unable to be reconciled by these algorithms that do not account for newly discovered interactions between the target DNA and RNP. For instance, a study by Malina *et al*. shows the presence of PAMs (Protospacer Adjacent Motifs) across the sequence to be inhibitory to Cas9 cleavage (Malina, et al., 2015). This underscores the need for additional experimental investigations into off-target effects and a robust algorithm to capture these effects for designing gRNAs, especially when applying CRISPR/Cas systems to non-model organisms.

Furthermore, the rapid discovery of novel, diverse Cas enzymes beyond spCas9 requires a flexible and robust algorithm that can accurately predict on- and off-target activities utilizing new CRISPR/Cas systems for genome editing. For instance, the discovery and use of another type of CRISPR class II endonuclease, *Acidaminococcus* sp. Cpf1 (asCpf1), has shown complementary activity to the Cas9 family with a T-rich PAM and “sticky-end” generation (Zetsche, et al., 2015). Based on the crystal structure of the asCpf1-RNA-DNA heterotriplex, it is possible to infer important nucleotides within the CRISPR-RNA (crRNA) for catalytic efficiency, and thus develop accurate targeting algorithms for genome editing (Kleinstiver, et al., 2016; Yamano, et al., 2016).

While previous research has focused on identifying highly active and unique target sites using on- and off-target algorithms to assist in gRNA design, targeting non-unique (repeated) sites with CRISPR tools may lead to some interesting future directions for experimentation (Prykhozhij, et al., 2015). This strategy, called multitargeting, has powerful metabolic engineering and synthetic biology applications, but has not yet been fully explored. For instance, repeated sequences may serve as sites where a single gRNA could induce knockouts across multiple orthologs of a gene, reducing the amount of heterologous machinery required to achieve the same effect. Furthermore, engineering consortia of organisms has been a burgeoning field in the past decade (Andrianantoandro, et al., 2006). CRISPR tools can potentially investigate and modify these consortia, which may not be achievable by conventional genetic manipulations. Software with the ability to implement these applications can pave the way for novel metabolic engineering and synthetic biology applications using CRISPR/Cas systems.

In this study, we developed a novel method, named CASPER (CRISPR associated software for pathway engineering and research) that implemented flexible algorithms to guide precise genome editing. Combining both experimental data and newly discovered biological principles, CASPER formulated an improved scoring method for enhanced prediction of on- and off-target activities. This prediction presents “relative” activities of a gRNA design that depend solely on target sequences, and hence are independent of experimental conditions (e.g., growth rate, Cas9 concentrations, and DNA supercoiling). These “relative” activities are different from “absolute” activities of a gRNA design that are condition-specific and must be determined experimentally (Farasat and Salis, 2016; Hsu, et al., 2013). Further, CASPER expanded novel applications of CRISPR/Cas systems in genome editing including multitargeting analysis (i.e. simultaneous multiple-site modification on a target genome with single gRNA requirement) and multipopulation analysis (i.e. gRNA design for genome editing of a consortium of organisms).

## 2. Methods

### On-target activity formulation for CRISPR gRNA design

On-target activity is defined as the binding affinity and subsequent nuclease activity of a particular gRNA and endonuclease complex to its matching DNA (or RNA) target sequence. Such activity depends on many factors that have been identified such as the state of DNA supercoils, genome size, Cas endonuclease concentration, and growth rate (Farasat and Salis, 2016). These variables change based on the organism and the experimental conditions and thus make it impractical to integrate into an algorithm designed for assessing a large set of non-model organisms and alternative Cas endonucleases. Such factors have the same effect on every target within a genome and therefore have no impact on the difference in relative activity between targets found on the same genome.

To predict the on-target activity of the complex, our developed CASPER method calculates the score S_C,P_ (equation 1) that is defined as follows:

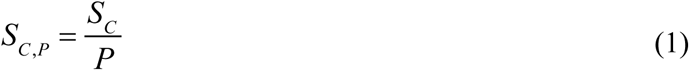

where S_C_ is the CRISPRscan (Moreno-Mateos, et al., 2015) score and P is the penalty score. In equation 1, S_C,P_ has a value between 0-100 that is assigned to a gRNA seed sequence, with higher values indicating higher predicted activity. The CRISPRscan score S_C_ (equation 2) is defined as follows:

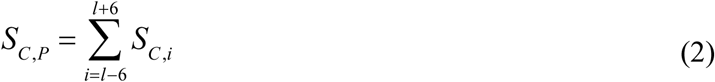

where l is the length of a seed sequence and PAM (see Supplementary Figure S1 for indexing illustration), and S_C,i_, is the score associated with the mono and dinucleotide features at position i (N_i_ or NN_i_) of the seed sequence that has been experimentally determined to be relevant (Moreno-Mateos, et al., 2015). Each score, S_C,i_, is the sum of the values of the features appearing at position i, and the sum of all positions’ scores (S_C_) is normalized with respect to the highest and lowest possible scores to obtain a normalized value from 0 to 100. The values of features at any given position can be seen in the CRISPRscan scoring table (Supplementary File 1).

The penalty score, P, (equation 3) is obtained by the combination of the PAM density score, s_ij_, (Supplementary Table S1) and the score S_G_ (equation 4):

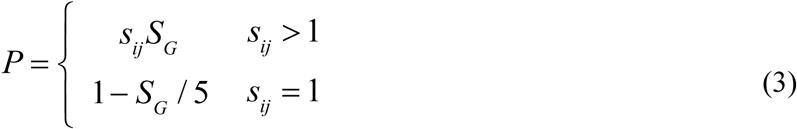

The score s_ij_ in equation 3 is determined by the number of PAMs present in a given seed sequence, where i is the number of PAMs in the forward sequence and j is the number of PAMs in the reverse complement of the sequence. The score s_ij_ can be looked up in the i^th^ column of the jth row in Supplementary Table S1, which is derived from experimental data (Malina, et al., 2015). Due to the length of the spCas9 PAM (3) and the length of a seed sequence (20), i + j < 8. In equation 3, the score S_G_, used to reinforce the importance of guanines and adenines to the stability/instability of the gRNA respectively, is formulated to account for the nucleotide composition of the seed sequence irrespective of position.

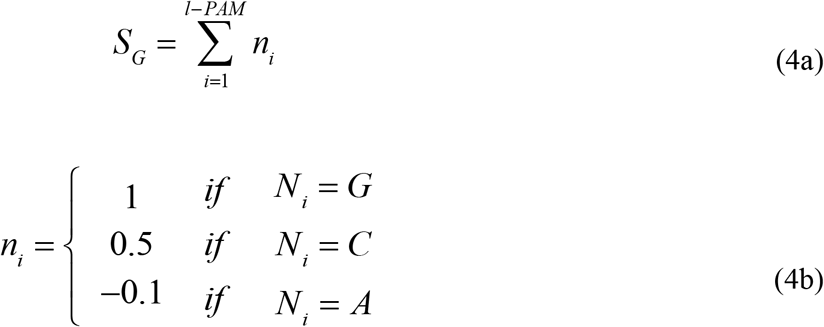

The values ni were derived to maximize correlation between scores and on-target experimental data (Moreno-Mateos, et al., 2015) further described in the Results. A visual summary of the process of obtaining an on-target score for a sequence is shown in Supplementary Fig. S1.

### Off-target formulation for CRISPR gRNA design

In contrast to the on-target activity assessment, the off-target activity is defined as the probability of a given gRNA sequence to interact with a non-matching sequence on the genome. Our developed CASPER method calculates the off-target score S_H,T,S_ as follows:

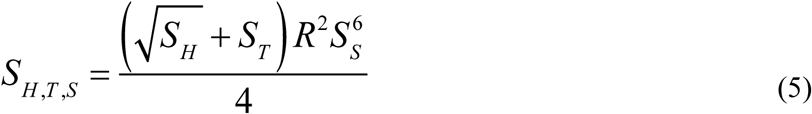

where the off-target score S_H,T,S_ lies between 0 and 1, with a higher value indicating higher probability of off-target activity, and is determined by combining four subscores S_H_, S_T_, S_S_, and R discussed below. The appropriate arrangement of these scores in equation 5 was determined by employing a genetic algorithm using the Pearson's coefficient between the output scores and experimental cleavage efficiency (Hsu, et al., 2013) as the fitness function. A more detailed description of the algorithm is presented in the Results.

The subscore S_H_ accounts for the types of nucleotide mismatches and their location on the seed sequence, and is defined as follows:

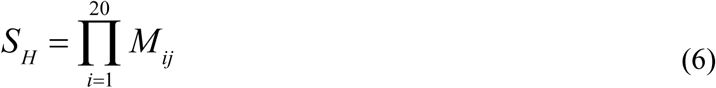

where M_ij_ is the element of the Hsu matrix derived from experimental data gathered in Hsu *et al*. (Hsu, et al., 2013) (Supplementary Table S2), i represents the index of the mismatch, and j corresponds to the identity of the mismatch (e.g. C with A).

The subscore S_T_ is derived from the inverse relationship of the proximity of the mismatch to the PAM:

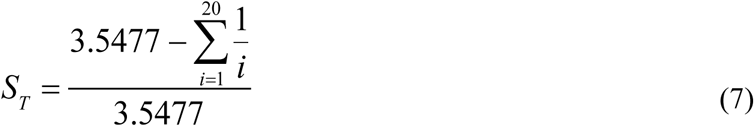

where S_T_ is valued from 0 to 1 with a higher score indicating a higher probability of off-target activity, and i is the index of the mismatch. This score was motivated by previous studies showing the farther a mismatch is from the PAM site, the less likely it is to interfere with activity (Anderson, et al., 2015). As a result, sequences with PAM distal mismatches are more likely to be sites of off-target activity. Summing the values for a mismatch at every location across the seed sequence gives a value of 3.5477, thus this number is used to normalize S_T_ as formulated in equation 7.

The subscore S_S_ also captures an inverse relationship of the proximity of the mismatch to the PAM but is formulated using a stepped scale:

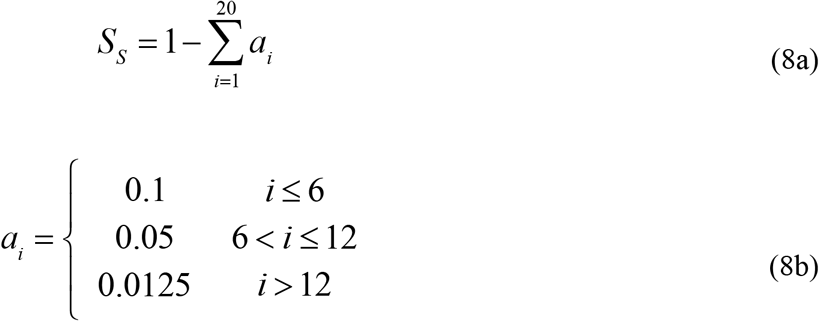

where i is the index of the mismatch, and a_i_ is defined by a step function. The step sizes were derived to agree with the previous experimental report where mismatches in regions closer to the PAM are more detrimental to activity (Hsu, et al., 2013).

Using the knowledge that some gRNAs are more stable/active than others, the on-target activity scores of the target sequence and the off-target sequence are assembled into a ratio, R:

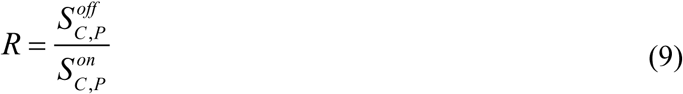

where S_C,P_^off^ and S_C,P_^on^ in equation 9 are the on-target scores for the target DNA sequences appearing at the undesirable and desirable sites of the genome, respectively. A ratio of R < 1 signifies that the sequence of interest (SOI) is more active compared to the potential off-target site, meaning less likelihood of the off-target site being hit compared to the SOI. A ratio R > 1 signifies the reverse, i.e. a highly active off-target site that has an increased likelihood of being hit compared to the SOI. The visual representation of gathering all these scores and combining them for a given two sequence comparison can be seen in Supplementary Figure S1.

### Multitargeting analysis

Multitargeting is the process of editing multiple sites simultaneously across the genome of an organism using a single gRNA. Instead of removing repeated gRNA target sequences, our developed CASPER method stores the data of repeated sequences for further analysis. To generate data of the non-unique seed sequences, CASPER references the set of sequences that appear more than once to a genome annotation file generated by either GenBank (http://www.ncbi.nlm.nih.gov/genbank/) or KEGG (genome.jp/kegg/). These sequences are then sorted based on the number of times they appear in an annotated region. Further analysis can be performed by investigating individual sequences in order to design gRNAs to target these repeated sequences.

### Multipopulation analysis

To analyze multiple genomes simultaneously for use in the study and genetic modification of a consortium, CASPER collects the data by identifying targets across a collection of genomes and performs on- and off-target activity analysis against the entire metagenome. This enables precise gRNA design for accurate genome editing within a consortium. Sequences shared across genomes can also be identified, thereby applying the concept of multitargeting to populations.

## 3. Results and Discussion

### 3.1 Development of CASPER algorithm for enhanced on-target activity prediction

In order to analyze sequences to be targeted by a gRNA/Cas complex, we developed an algorithm for CASPER to predict the gRNA design for the on-target activity (equation 1). It was formulated to incorporate three guiding principles: i) the CRISPRscan features experimentally identified to be present in highly active guide sequences (Moreno-Mateos, et al., 2015), ii) the density of the PAM in question across the guide sequence (Malina, et al., 2015), and iii) the propensity for guanines (and to a lesser extent cytosines) over adenines in the gRNA sequence which has been shown to be a factor in gRNA stability (Doench, et al., 2014; Wang, et al., 2014). To formulate a combination of these three principles, we combined the normalized CRISPRscan score, S_C_ (equation 2) and divided it by a penalty score, P (equation 3). The structure of P is determined by whether or not the number of PAMs in the sequence is inhibitory (s_ij_ > 1). If so, the value is multiplied by the score S_G_; otherwise, it is disregarded and P purely accounts for the nucleotide content of the sequence (the S_G_ score). While the values of s_ij_ are derived directly from experimental data (Malina, et al., 2015), the score S_G_ needs to be constructed *de novo.*

Since previous studies showed guanines were favorable and adenines unfavorable to gRNA stability (Doench, et al., 2014; Wang, et al., 2014), we decided to value guanine most favorably (positive value) and adenine least favorably (negative value). The values assigned to guanine, adenine, and cytosine (equation 4b) were obtained by varying their values between 0 to 1 (with a step of 0.1) and then evaluating which combination of values resulted in the optimal Pearson coefficient between experimental data from Moreno-Mateos *et al*. (Moreno-Mateos, et al., 2015) and the score, S_C,P_.

We applied CASPER's on-target activity algorithm to predict the on-target activities of gRNAs from the experimental study by Moreno-Mateos et al. (Moreno-Mateos, et al., 2015). This study investigated indel frequency from a pool of 1280 gRNAs in zebrafish single cell embryos at 9-hour post fertilization. The CASPER method provided a minor 2.4% improvement in the R^2^ value as compared to the CRISPRscan model (Figure 1A) by taking into account PAM density and position-independent nucleotide content. Such a minor variation underscores the robustness of the CRISPRscan model. This improvement is statistically insignificant from the CRISPRscan model as determined by an F-test between the two data sets (p<0.05). However, CASPER successfully identified 3 of the 4 largest CRISPRscan score outliers in an experimental dataset of 25 gRNAs (Figure 1A).

**Fig. 1.**
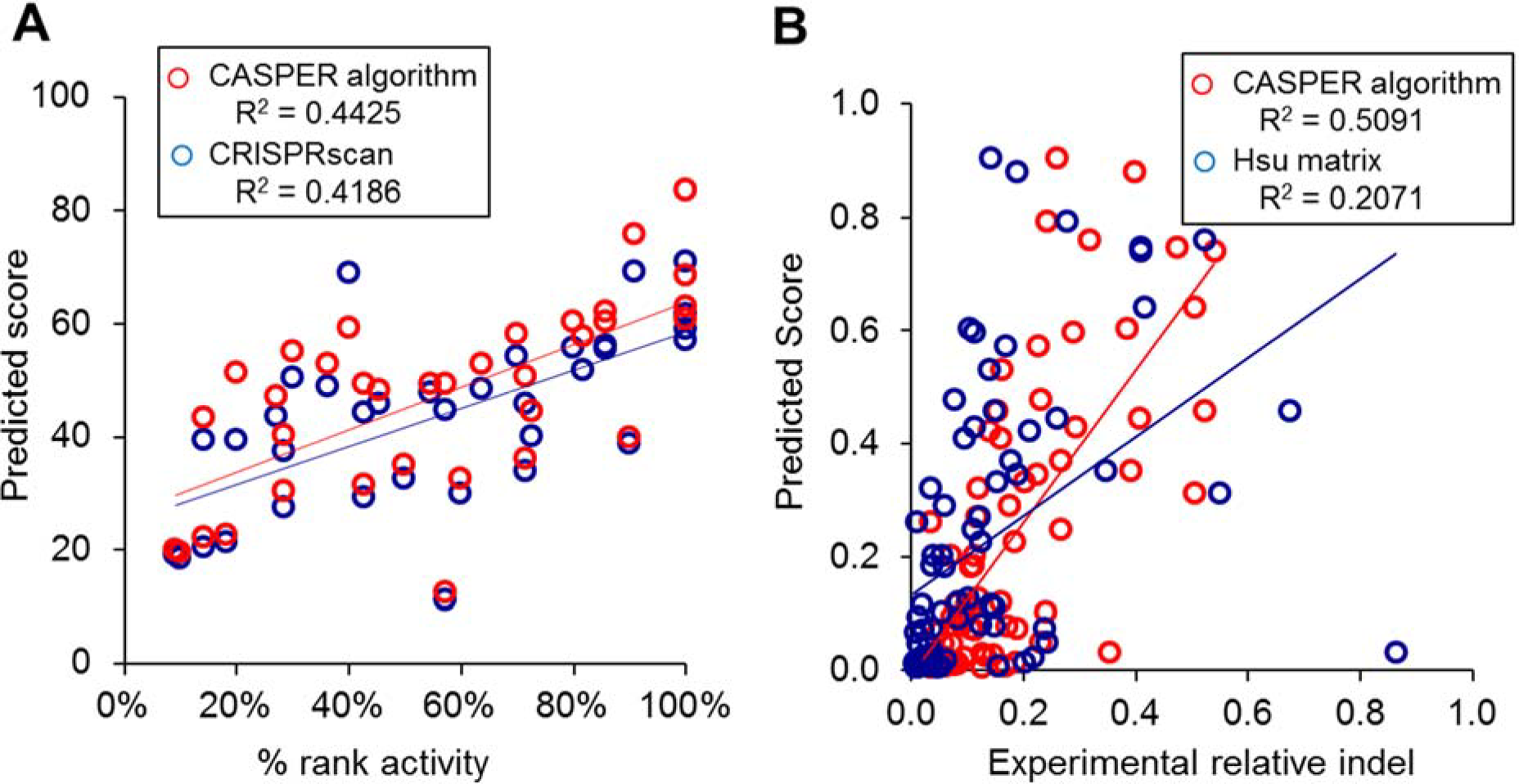
**(A)** Comparison of CASPER on-target algorithm with CRISPRscan against experimental data obtained from Moreno-Mateos *et al*. (Moreno-Mateos, et al., 2015). The % rank activity (on-target) is obtained by ranking the activity of a sequence with respect to the rest of other sequences in the sample set. **(B)** Comparison of CASPER off-target algorithm with Hsu matrix method against experimental data obtained from Hsu *et al*. (off-target) (Hsu, et al., 2013). The experimental relative indel (off-target) is determined by dividing the absolute indel % of the off-target site by the indel % of the targeted site. For example, a gRNA that has a 25.0% indel at the targeted site and 1.0% at the off-target site gives a 4% (or 0.04) relative indel.

CASPER's on-target activity algorithm can be refined with subsequent data sets or further studies on spCas9's underlying mechanism of action. Additional on-target data sets generated by future studies will allow CASPER to develop parameters to assess gRNA activity in novel CRISPR/Cas systems (i.e. diverse organisms and Cas enzymes). For instance, CASPER's on-target activity algorithm can be applied directly to asCpf1 as soon as experimental data of on-target activity becomes available. In the meantime, to evaluate asCpf1 and other non-canonical endonucleases, CASPER focuses on the principles governing PAM density and guanine/adenine content to predict on-target activity.

### 3.2 Development of CASPER algorithm for enhanced off-target activity prediction

The objective of off-target analysis is to determine the sequences most likely to exhibit activity despite not being an identical match to the designed gRNA. Many factors contribute to off-target activity, including the number and location of mismatches, the number of times the PAM appears across the genome, and the concentration of the endonuclease. Only those variables relating to sequence identity are considered here, as other factors are considered to be either relatively constant (e.g. Cas9 concentration) or entirely unpredictable (e.g. state of DNA supercoils) when comparing one sequence relative to another. To perform such an analysis, an algorithm was developed for CASPER to compare all potential target sites in the genome to the SOI. For each comparison, mismatches between the two sequences are identified. If the total number of mismatches across the two sequences exceeds 4 the pairing is given a score of 0, signifying there should be no appreciable activity at the off-target sequence in question. The mismatches are then scored according to equations 6, 7, and 8a. In addition, the on-target scores for the two sequences are obtained and the ratio (equation 9) is also incorporated into a final score, S_H,T,S_ (equation 5). This scoring method improved prediction capabilities by 30.2% in the R^2^ value (Figure 1B) over the canonical study (Hsu, et al., 2013).

The enhanced correlation to experimental values could be attributed to the combination of incorporating experimental data (subscore S_H_) (Hsu, et al., 2013) and guiding principles derived from the CRISPR RNP complex's mechanisms of action (subscores S_T_, S_S_, R). A genetic algorithm was employed to determine the optimum arrangement of the subscores (S_H_, S_T_, S_S_, and R) to create a single final score, S_H,T,S_, and to determine if any subscore was redundant. The result from the combinations of these scores were compared to experimental values given in Hsu *et al*. (Hsu, et al., 2013) and the resulting correlation (R^2^) was used as the fitness parameter for the genetic algorithm. The algorithm was given free range over modifying the coefficients and exponents for each of the subscores, as well as the general form of the equation, i.e. whether the subscores were added/subtracted or multiplied/divided to each other. The algorithm was initialized with 100 possibilities and the fittest “parents” were chosen for crossover to create a new generation. Each generation spawned 100 children. By running the algorithm for 1,000 generations, the final format present in equation 5 was achieved. To confirm the algorithm was not over trained on the experimental data set provided, a separate set of data (Hsu, et al., 2013) was used for determining the correlation of the algorithm's output with experimental off-target activity data (Figure 1B).

Off-target guiding principles of asCpf1 are similar to those of Cas9 in that the PAM proximal nucleotides are important for binding and enzyme activity (Zetsche, et al., 2015). asCpf1 off-target identification was simplified to the use of equations 7–9 as off-target data to generate a matrix for equation 6 were not available at the time of this study. Using the recently solved crystal structure of the asCpf1-RNA-DNA heterotriplex (Yamano, et al., 2016), parameters of equation 8 were modified to give equation 10:

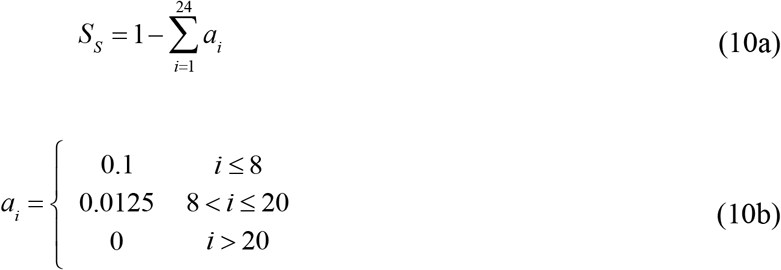

As in equation 8b, the values for ai in 10b are calculated such that a completely mismatched sequence is given a score (S_S_) of 0. This equation accounts for experimental observations that the most significant nucleotides in the seed sequence for asCpf1 are the 8 nucleotides proximal to the PAM. In the absence of experimental data, to perform off-target activity analysis on non-canonical CRISPR endonucleases, such as asCpf1, CASPER subscores can be used independently to evaluate off-target activities.

The 30.2% increase in R^2^ value was evaluated for statistical significance by an F-test using the scores from the Hsu matrix. It was confirmed that the CASPER algorithm represents a statistically significant improvement (p<0.05). In more measurable terms, the off-target activity of roughly 3 out of 10 sequences evaluated will be falsely predicted by the Hsu matrix that CASPER can reconcile.

The newly developed CASPER off-target algorithm can be quickly adapted to other systems seamlessly by training the algorithm against a set of experimental data and accounting for biophysical properties of the CRISPR RNP complex to minimize experimental bias. CASPER therefore provides a foundation for establishing *in silico* off-target analysis for a variety of organism and endonuclease combinations, especially when dealing with non-model organisms.

### 3.3 Development of multitargeting analysis

While sequences repeated in a genome have been traditionally discarded due to their inherent lack of specificity, we developed CASPER to exploit these sequences as potential gRNAs that are capable of targeting multiple sites across a genome with useful applications in synthetic biology and metabolic engineering.

To demonstrate CASPER for multitargeting analysis, we compiled data for 34 genomes of model and non-model organisms with three different endonucleases (spCas9, asCpf1, and spCas9-VRER) to gain perspective on the number of target sites that appear across each genome (Figure 2). Genomes with a low GC content such as *Clostridium beijerinckii* (31.0%) and *C. saccharobutylicum* (29.4%) have a much greater number of target sites for asCpf1 (PAM: TTTN) than that of spCas9 (PAM: NGG). The spCas9-VRER variant (PAM: NGCG) in particular have only a couple thousand sequences appear for the mentioned organisms, a two order of magnitude difference compared to asCpf1. In addition, the greater PAM length of the VRER variant, which one would assume to appear less frequently than the canonical NGG, does indeed present less target sites across all the organisms investigated. These results show that some endonucleases are more useful than others depending on the genome and application. While this paper only presents a representative sample of non-model organisms, the algorithm is capable of analyzing any desired combination of organisms and endonucleases.

**Fig. 2.**
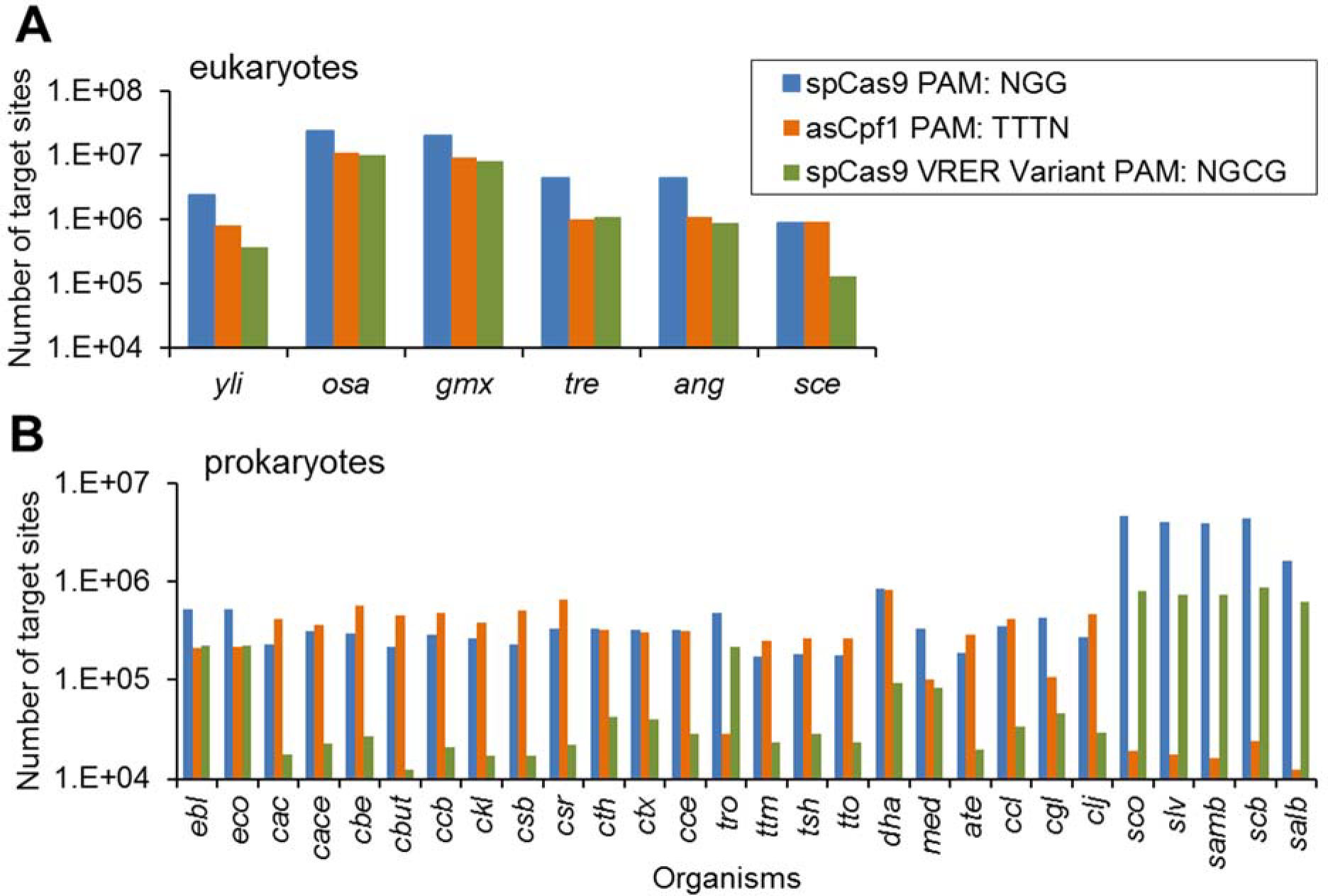
Appearances of PAM motifs for three common Cas endonucleases (spCas9, asCpf1, and spCas9-VRER variant) across a select number of non-model organisms including **(A)** eukaryotes and **(B)** prokaryotes. Organism abbreviations: *yli: Yarrowia lipolytica, ang: Aspergillus niger, sce: Saccharomyces cerevisiae gmx: Glycine max (soybean) osa: Oryza sativa Japonica (rice) tre: Trichoderma reesei, cac: Clostridium acetobutylicum, cace: Clostridium aceticum, cbe: Clostridium beijerinckii, cbut: Clostridium butylicum, ccb: Clostridium cellulovorans, ckl: Clostridium kluyveri, csb: Clostridium saccharobutylicum, csr: Clostridium saccharoperbuytlacetonium, cth: Clostridium thermocellum (ATCC 27405), ctx: Clostridium thermocellum (DSM 1313), cce: Clostridium cellulolyticum, tro: Thermomicrobium roseum, ttm: Thermoanaerobacterium thermosaccharolyticum (DSM 571), tsh: Thermoanerobacterium saccharolyticum, tto: Thermoanaerobacterium thermosaccharolyticum (M0795), dha: Debaryomyces hanseii, med: Megasphaera elsdenii, ate: Caldicellulosiruptor bescii, ccl: Clostridium clariflavum, cgl: Cornyebacterium glutamicum, clj: Clostridium ljungdahlii, sco: Streptomyces coelicolor, slv: Streptomyces lividans, samb: Streptomyces ambofaciens, scb: Streptomyces scabiei, salb: Streptomyces albus.*

To understand where these repeated sequences appear, we mapped their locations on the annotated genomes. Figure 3 reveals that many of these sequences appeared in annotated regions, but a significant number also fell in unannotated ones, particularly sequences targeted by asCpf1. This discovery is valuable in that it helps reveal regions of the genome that may be related and subsequently probed through the precise CRISPR genome editing tool. Further analysis into non-unique targets in unannotated regions revealed a significant number of sequences in regions of completely unknown function. Sequences appearing in unannotated regions are particularly useful for two reasons. First, designing gRNAs to target sequences that appear in unannotated regions may reveal information about whether these regions are functional. Second, these sequences can be used for inserting multiple copies of genetic cargo with a reduced risk of unintentional disruption of cellular function. With multitargeting analysis, CASPER facilitates the investigation and manipulation of unannotated regions with the CRISPR/Cas system for rapid genome editing.

**Fig. 3.**
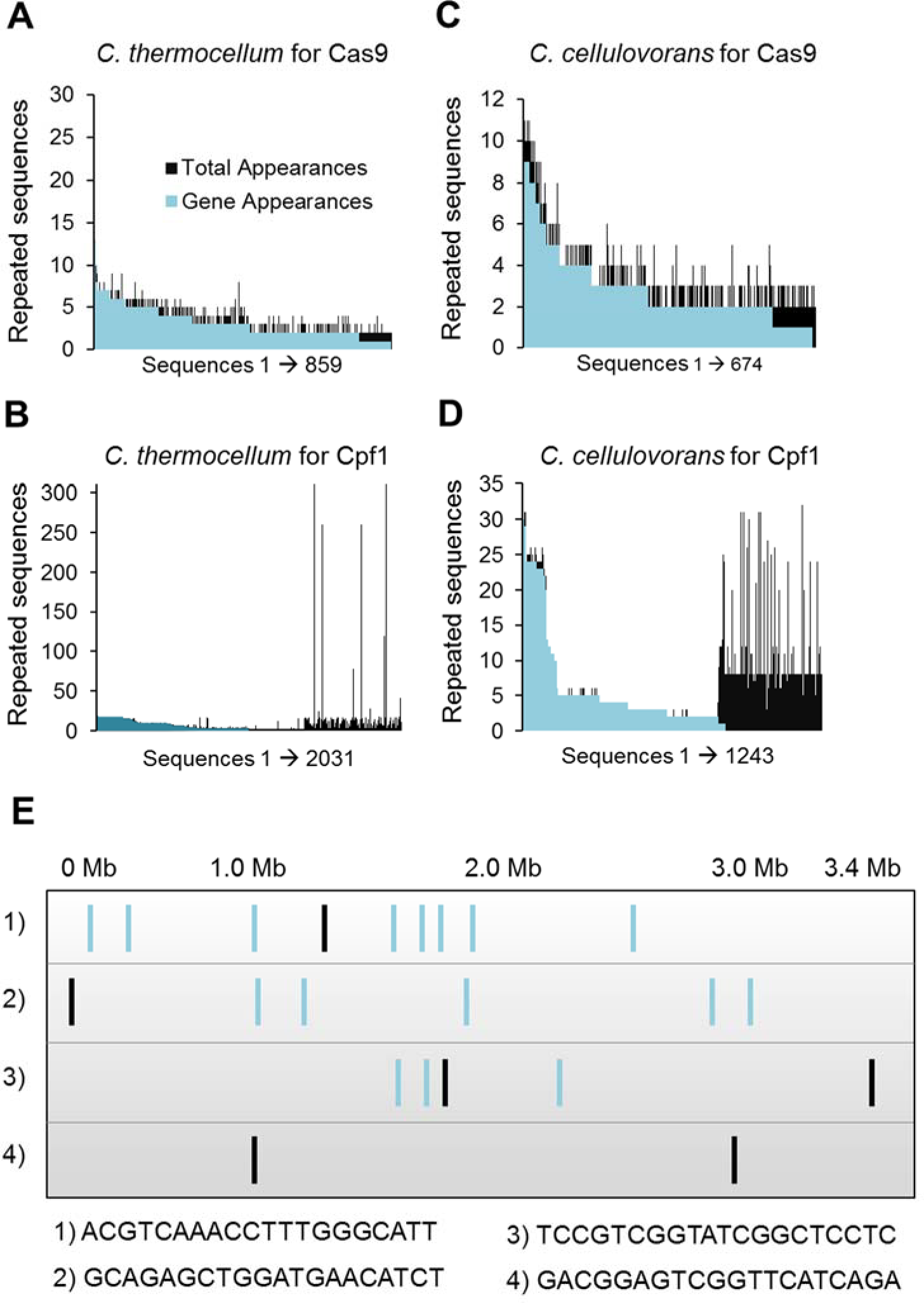
Multitargeting analysis for **(A)** *C. thermocellum* using spCas9, **(B)** *C. cellulovorans* using spCas9, **(C)** *C. thermocellum* using asCpf1, and **(D)** *C. cellulovorans* using asCpf1. **(E)** The distribution across the genome of *C. thermocellum* using a set of 4 representative non-unique sequences. Green lines indicate appearance within a gene sequence while black lines represent appearances in un-annotated regions.

To discover cellular processes that lend themselves to multitargeting, we looked further into the identity of the annotated regions in which these sequences were appearing. A cursory examination of the repeated sequences across the genomes revealed that transposable elements appear frequently across all organisms and can provide a powerful platform for integration of multiple copies of genes or entire operons (Figure 4). Across the species investigated, between half and two-thirds of the non-unique sequences found on annotated regions were located on a transposon related feature. The promise of harnessing transposons for genome manipulation has been well documented by the utility of the Sleeping Beauty transposon system (e.g. SB10) (Geurts, et al., 2003; Ivics, et al., 1997). In addition to transposable elements, regions labeled as hypothetical proteins are also common sites to find repeated sequences (Figure 4). Targets within hypothetical protein regions may be useful in a similar manner to those appearing in unannotated regions because the function of these cryptic regions can be systematically investigated.

**Fig. 4.**
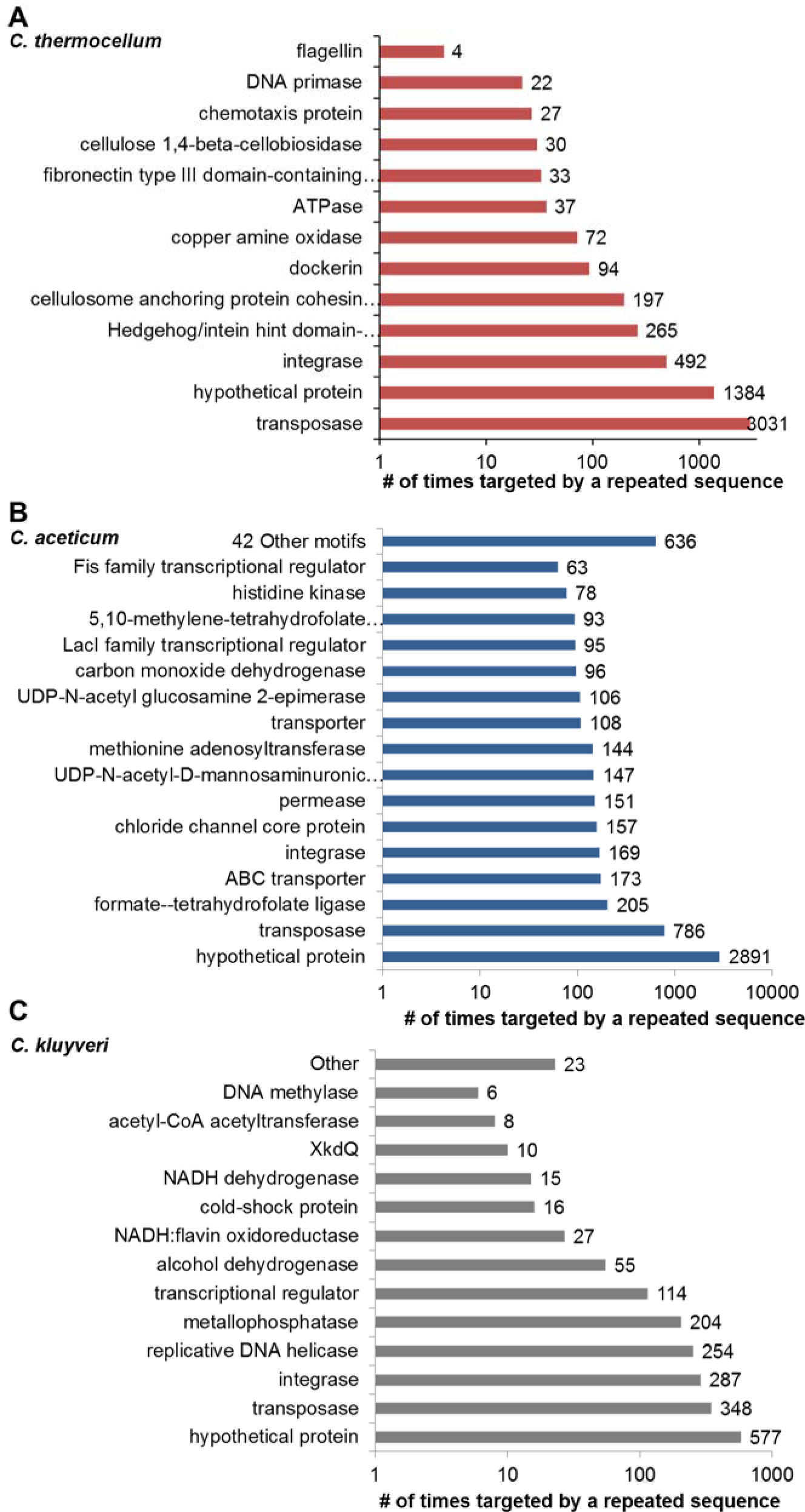
Annotated cellular processes targeted by repeated sequences for **(A)** *C. thermocellum,* **(B)** *C. acetobutylicum,* and **(C)** *C. kluyveri*.

Of particular interest are also the repeated sequences that appear on gene loci. Figure 4 details the common motifs targeted by non-unique sequences across the genomes of *C. thermocellum, C. aceticum,* and *C. kluyveri.* Interestingly, elements such as the cellulosome anchoring protein of *C. thermocellum*, a defining feature of the organism, is targeted 197 times by repeated sequences, opening the possibility of interrogating this structure with a select number of gRNAs. In *S. cerevisiae,* hexose transporters (416 times) and heat-shock proteins (111 times) are cellular processes that also may be targeted simultaneously by a single gRNA and are of particular interest for metabolic engineers looking to probe the sugar metabolism and environmental sensitivity and adaptation in this organism. In general, CASPER is capable of performing multitargeting analysis in any organism and with any endonuclease desired.

### 3.4 Development of multipopulation analysis

When editing a consortium of organisms with a CRISPR/Cas system, one must be mindful of the existence of similar or even shared sequences among genomes. Horizontal gene transfer of a plasmid containing the CRISPR/Cas system can exhibit activity in other species’ genomes within a consortium. Additionally, direct transformation of the RNP into a consortium may result in unintended off-target activity in multiple species. It is therefore important to screen all genomes in a consortium for potential off-targets. We have developed the CASPER off-target algorithm to check for off-target sites across genomes in the consortium, thus minimizing unintended activity not just within the targeted organism but the consortium as a whole. In addition, CASPER can identify repeated sequences across the genomes that may lend themselves to multitargeting. This enables the identification of potential sites where multiple organisms in the consortium can be edited with the same gRNA.

To examine more closely the opportunities for multitargeting in a consortium, we ran CASPER against pairs of *Clostridial* species. Figure 5A-B details the number of sequences available for such targeting between the pairing of *C. thermocellum* and a selection of thermophilic species, as well as the pairing of *C. cellulolyticum* with other mesophiles. Additionally, CASPER is capable of performing an analysis on multiple organisms to identify repeated sequences shared across them. For instance, we applied CASPER to identify repeated sites across *C. kluyveri, C. cellulolyticum*, and *C. aceticum* (Figure 5C-D). This consortium was designed to utilize either biomass or a hydrogen/carbon dioxide feed for production of industrially relevant long-chain organic acids. This example demonstrates how CASPER can be used to identify off-targets and multitargeting analysis for genome editing of a consortium.

**Fig. 5.**
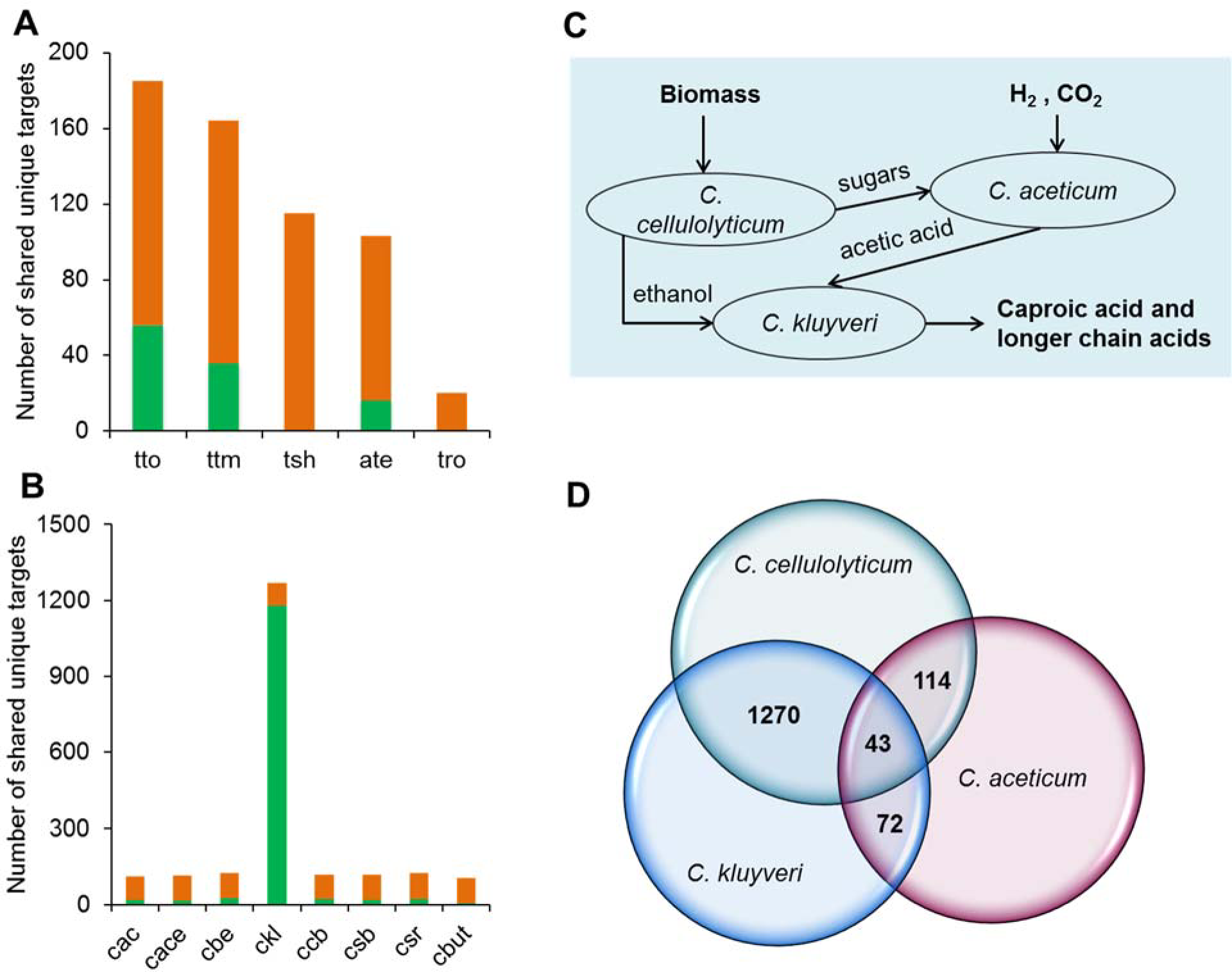
Comparison of the number of shared targets between **(A)** *C. thermocellum* and other thermophiles and **B)** *C. cellulolyticum* and other mesophiles. The green portion of the bar represents the number of sequences that appear in both genomes but are unique within each genome. Orange portion of the bar represents the number of sequences that appear in both genomes and is repeated within one or both genomes being compared. **(C)** A hypothetical synthetic consortium of three different *Clostridial* species with synergistic metabolisms. The Venn diagram represents a three-way comparison of these organisms with the numbers of sequences shared among the genomes shown at the intersections.

The number of sequences that may be used to target multiple organisms is quite small when compared to the size of the genomes. Thus we postulate that the usefulness of multitargeting across organisms lies in niche insertions or deletions, such as those targeting a gene with high sequence similarity across the organisms in question. The opportunities for such manipulation will vary drastically depending on the identity of a consortium. Further, off-target analysis for a consortium will prove crucial in preventing unintended activity. Overall, CASPER's algorithms can facilitate the use of CRISPR tools to edit target genomes within a heterogeneous consortium.

## 4. Conclusion

The development of CASPER's flexible algorithms for analyzing on- and off-target activity in any organism with any Cas enzyme broadens the utility of CRISPR tools for genome editing of industrially relevant and non-model organisms. CASPER's multitargeting analysis facilitates simultaneous genetic manipulation of multiple loci using a single gRNA with novel potential applications including the investigation of large complex systems (e.g. the cellulosome of *C. thermocellum*) and unannotated genome regions. Further, CASPER's multipopulation analysis provides the ability to investigate microbial consortia and apply the CRISPR tools to perform genetic manipulations on single or multiple organisms within the consortia. We envision CASPER will assist the progress of CRISPR genome editing for metabolic engineering and synthetic biology applications.

## Acknowledgements

The authors would like to thank Trinh lab's members for useful comments.

